# Deciphering sepsis molecular subtypes using large-scale data to identify subtype-specific drug repurposing

**DOI:** 10.64898/2026.03.28.714506

**Authors:** Leslie A. Smith, Beulah Augustin, Vinitha Jacob, Lauren P. Black, Andrew Bertrand, Charlotte Hopson, Emilio Cagmat, Susmita Datta, Srinivasa T. Reddy, Faheem W. Guirgis, Kiley Graim

**Affiliations:** University of Florida, Herbert Wertheim College of Engineering, Department of Computer and Information Science and Engineering, Gainesville, FL; University of Florida, College of Medicine, Department of Emergency Medicine, Gainesville, FL; University of Michigan, School of Medicine, Department of Emergency Medicine, Ann Arbor, Michigan; Northwestern University, Feinberg School of Medicine, Department of Emergency Medicine, Chicago, IL; University of Florida, College of Public Health and Health Professions, Department of Biostatistics, Gainesville, Florida; University of California Los Angeles, David Geffen School of Medicine, Department of Medicine, Los Angeles California; University of Florida Health Cancer Center, Gainesville, FL; University of Florida Genetics Institute, Gainesville, FL; Mayo Clinic Jacksonville, Jacksonville, FL

**Keywords:** Sepsis, Drug Repurposing, Transcriptomics

## Abstract

Sepsis is a life-threatening dysregulated response to infection, the heterogeneity of which precludes effective precision medicine. Deciphering this heterogeneity requires substantially larger molecular datasets of sepsis patients. Here, we create a transcriptomic atlas of publicly available adult sepsis data. In total, we harmonized data from 3,713 samples across 28 datasets, including 2,251 unique sepsis patients. On this data, we performed molecular subtyping and identified potential subtype-specific drug repurposing opportunities. We identified 4 molecular subtypes of sepsis (C1 – C4) based on expression differences in immune- and lipid-related genes. Pathway analysis of gene signatures unique to each molecular subtype revealed patterns of immune exhaustion and metabolic dysregulation in C1, suggesting potential benefit from corticosteroid treatment. C2 expression patterns were often anti-correlated with those of C1, and C2 had the youngest patient population and the lowest mortality. C3 was enriched for inflammatory and cellular stress pathways, while the highest mortality subtype, C4, showed evidence of immunosuppression and metabolic reprogramming. Gene and pathway-level analysis of our molecular subtypes statistically correlated with results from analysis of 28-day mortality, with the best (C2) and worst subtypes (C4) exhibiting similar molecular dysregulation as survivors and non-survivors, respectively, suggesting that targeted subtype-specific approaches could improve sepsis management. Using a large-scale pharmacogenomics database, ClinPGx, we then evaluated potential targeted therapies for each subtype and assessed the potential clinical impacts of these drugs. We identified several potential subtype-specific candidate therapies, including possible responsiveness to Methylene Blue therapy for patients from our highest mortality subtype, C4. Notably, our drug repurposing analysis revealed a significant representation of anti-inflammatory monoclonal antibody therapies across molecular subtypes. The anti-correlated signatures in C1 and C2 suggest that monoclonal antibody therapies may only be effective for patients in one of the subtypes, which may explain why prior clinical trials have been unsuccessful. Altogether, our detailed molecular subtyping and drug repurposing analysis identified subtype-specific potential drug targets, with implications for future precision medicine for sepsis.

## Introduction

Sepsis is a potentially deadly dysregulated response to infection^1,2^ with an increasing global burden^3^. In the United States, sepsis costs more than $24 billion annually and is the most common cause of in-hospital deaths^3–5^. Changes in diagnostic coding, an aging population, increases in chronic illnesses and immunocompromising comorbidities, and the rise of antimicrobial resistance are hypothesized to drive increases in sepsis incidence^6–9^. The highly heterogeneous nature of sepsis has stymied success in clinical trials for over 30 years^10–15^. Transcriptomic biomarkers have shown increasing utility in aiding treatment decisions in several diseases, enabling precision medicine approaches^16–18^. In sepsis, use of transcriptomic biomarkers in these tasks provides a promising approach because gene expression changes accurately reflect disease states during disease progression and provide a better picture of the patient’s evolving immune landscape^9,19–21^.

Unsupervised clustering has been successfully used to identify clinically relevant subtypes driving differences in disease progression, mortality and treatment efficacy across various conditions^22–27^. Unsupervised clustering of omics data has been particularly insightful in understanding the biological mechanisms driving disease heterogeneity^16,17,28–31^. However, in sepsis, subtypes identified in previous studies are often limited by the availability of data, size of the patient cohort, clustering algorithm used for analysis, and the biology of interest^32^. Here, we create a 28-study sepsis atlas from which we identify robust, reproducible sepsis molecular subtypes via unsupervised clustering of transcriptomic data in a large cohort of samples to decipher the molecular signals underlying sepsis heterogeneity, contextualize sepsis variation and identify possible molecular subtype-specific drug targets.

## Methods

### Creating the sepsis transcriptomic data atlas

This is a secondary analysis of transcriptomic data from 28 sepsis and septic shock studies published between 2009 and 2024. We surveyed three genomic databases (refine.bio, GEO, and SRA) for studies with samples that met our predefined inclusion criteria (Table 1). Samples were from adult patients meeting sepsis or septic shock criteria with whole blood bulk mRNA expression data available and sampled within 48 hours of sepsis diagnosis. Patients with parasitic infections or non-bacterial sepsis were excluded.

**Table 1.** Study inclusion criteria. Inclusion criteria and exclusion criteria used for study screening across databases.

| Inclusion Criteria |
| --- |
| Adult samples only (no pediatric) |
| Meeting sepsis or septic/shock criteria |
| Bulk RNA expression data |
| Whole blood samples only |
| Blood drawn within 48 hours of sepsis recognition |
| If multiple time points for samples, only earliest is included for analysis |
| Exclusion Criteria |
| Pediatric |
| Immunocompromised |
| Death within 12 hours of hospital admission |
| COVID-19 diagnosis |
| Parasitic infections |
| Sepsis melioidosis |

Raw sequencing reads from study datasets were processed separately before harmonization of all data (Figure 1 and Material and Methods Supplement). Following individual dataset processing, we combined the 3,713 samples from all studies into one harmonized transcriptomic dataset, including healthy controls and diseases other than sepsis. We corrected for batch effects using all samples in this merged dataset, then removed all samples that were not sepsis or septic shock. For patients with samples taken at multiple timepoints, only the first timepoint was retained. After data harmonization and batch correction using ComBat (sva version 3.48)^33^ the sepsis data atlas included 2,251 samples and 10,150 genes. Both the 3,713 samples and 2,251 sample datasets are provided. Metadata was manually curated for each sample included in the data atlas. To do this, we created a joint metadata file with all samples (Supplemental Table 1). Columns existing in all or most studies (e.g. age, sex, survival) were integrated into a single merged column and inserted into the joint metadata file (see Table 2).

**Figure 1.**
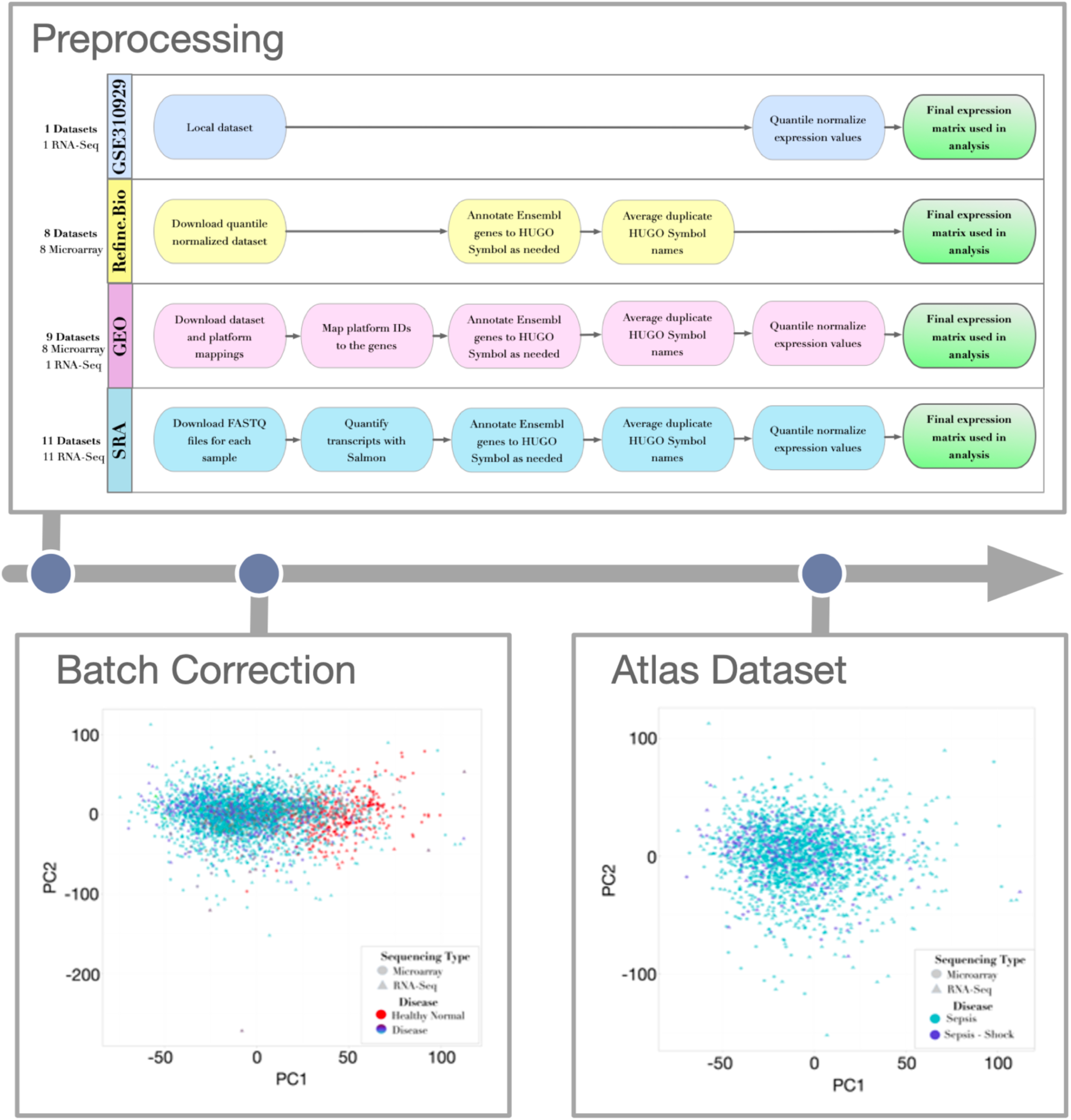
Study Pipeline Overview. We processed 28 studies from 4 sources. After preprocessing each dataset, all datasets are combined, and all samples are included for batch correction. Following batch correction, samples classified as having sepsis and septic shock are clustered and analyzed.

**Figure 2.**
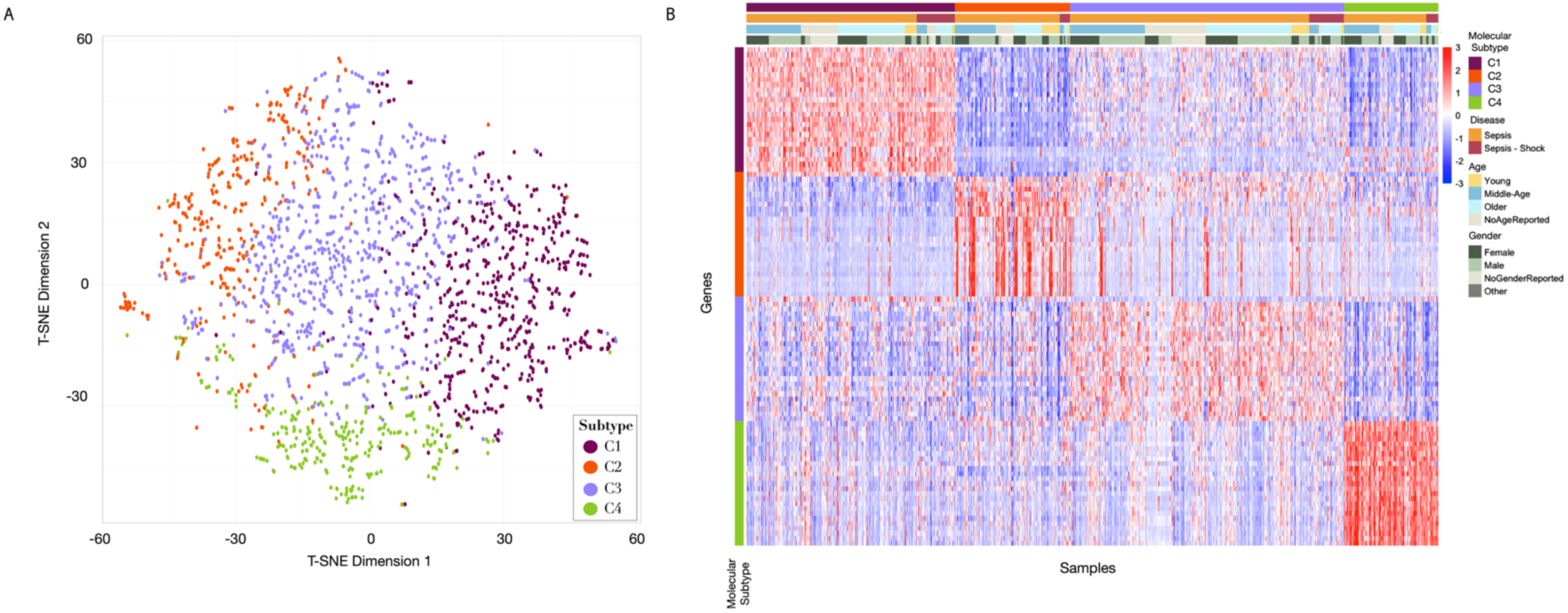
t-SNE and Heatmap of C1-C4 Identified Molecular Subtypes. (A) t-SNE plot depicting the four molecular subtypes identified by K-Means clustering. (B) Heatmap showing the gene expression of the top 25 genes with the largest log fold change for each molecular subtype. Rows depict gene values and are sorted by molecular subtype, columns are samples and are sorted first by molecular subtype, then by disease (sepsis vs septic shock), then by age, and finally by gender. The heatmap reveals distinct gene upregulation across molecular subtypes.

**Table 2.** Summary of metadata information available for all included studies. Missing values are notated, and all proportions are calculated relative to the non-missing cohort for that variable *Age: 20 samples with only 20-year age ranges provided assumed to be in the middle of those ranges (all from Dataset GSE33118) **Gender: 2 samples labeled as “H” assumed to be missing (both from Dataset GSE95233). ***Race: 1 Native American combined with 21 Other, 3 Unknown considered to be Missing

| Dataset ID | First Author | Sample Size | Age* | Gender – Male** | Race*** | Reported Sepsis, Shock | Survival |
| --- | --- | --- | --- | --- | --- | --- | --- |
| GSE65682 | Scicluna et al | 479 | 63.0 [53.0, 71.0] | 272 (57%) | - | 479 (100%), 0 (0%) | 365 (76%) |
| GSE185263 | Baghela et al | 348 | 61.0 [43.8, 72.3] | 202 (58%) | - | 348 (100%), 0 (0%) | 293 (85%)<br>(3 missing) |
| Faheem | Guirgis et al, Augustin et al | 193 | 62.2 [56.1, 72.0] | 108 (56%) | Black: 100 (52%)<br>Asian: 1 (<1%)<br>White: 87 (45%)<br>Other: 5 (3%) | 94 (49%), 99 (51%) | 169 (88%) |
| GSE236892 | Neyton et al | 189 | 65.0 [56.0, 77.0] | 113 (60%) | - | 189 (100%), 0 (0%) | 123 (65%) |
| GSE134347 | Scicluna et al | 156 | 61.5 [49.8, 70.3] | 98 (63%) | - | 156 (100%), 0 (0%) | - |
| GSE236713 | Szakmany et al | 125 | - | 65 (52%) | - | 125 (100%), 0 (0%) | 95 (76%) |
| GSE189400 | Kalantar et al | 121 | 67.0 [56.0, 78.0] | 75 (62%) | Black: 20 (17%)<br>Asian: 36 (31%)<br>White: 45 (38%)<br>Other: 17 (14%)<br>(3 missing) | 121 (100%), 0 (0%) | 86 (71%) |
| GSE131761 | Martinez-Paz et al | 81 | 75.0 [66.0, 80.0] | 48 (59%) | - | 0 (0%), 81 (100%) | 42 (52%) |
| GSE74224 | McHugh et al | 74 | - | - | - | 74 (100%), 0 (0%) | - |
| GSE66890 | Kangelaris et al | 54 | 65.0 [49.3, 77.8] | 31 (57%) | - | 23 (43%), 31 (57%) | - |
| GSE95233 | Venet et al | 49 | 66.0 [54.0, 75.0] | 30 (64%)<br>(2 missing) | - | 0 (0%), 49 (100%) | 33 (67%) |
| GSE32707 | Dolinay et al | 48 | - | - | - | 48 (100%), 0 (0%) | - |
| GSE154918 | Herwanto et al | 39 | - | 19 (49%) | - | 20 (51%), 19 (49%) | 36 (92%) |
| GSE216902 | Chen et al | 37 | - | - | - | 37 (100%), 0 (0%) | - |
| GSE63311 | Pena et al | 37 | - | - | - | 37 (100%), 0 (0%) | - |
| GSE57065 | Cazalis et al | 28 | 62.0 [54.3, 76.0] | 19 (68%) | - | 0 (0%), 28 (100%) | - |
| GSE13015 | Pankla et al | 24 | 55.0 [50.0, 66.8] | 10 (42%) | Asian: 24 (100%) | 14 (58%), 10 (42%) | 17 (71%) |
| GSE137340 | - | 21 | 42.0 [30.0, 63.0] | 14 (67%) | - | 15 (71%), 6 (29%) | 18 (95%)<br>(2 missing) |
| SRP198776 | Braga et al | 21 | 73.0 [62.0, 80.0] | 14 (67%) | - | 0 (0%), 21 (100%) | 16 (76%) |
| GSE232753 | Choi et al | 20 | - | - | - | 20 (100%), 0 (0%) | - |
| GSE33118 | - | 20 | 50.0 [50.0, 60.0] | 12 (60%) | - | 0 (0%), 20 (100%) | 10 (50%) |
| GSE69063 | - | 19 | - | - | - | 19 (100%), 0 (0%) | - |
| GSE196117 | Karakike et al | 18 | 67.0 [55.5, 72.5] | 11 (61%) | - | 18 (100%), 0 (0%) | - |
| GSE199816 | Ito et al | 18 | - | - | - | 18 (100%), 0 (0%) | - |
| GSE100159 | Rinchai et al | 17 | - | - | - | 17 (100%), 0 (0%) | - |
| GSE211210 | Yang et al | 5 | - | - | - | 5 (100%), 0 (0%) | - |
| GSE222393 | An et al | 5 | 66.0 [55.0, 72.0] | 4 (80%) | - | 5 (100%), 0 (0%) | 4 (80%) |
| GSE232404 | Ma et al | 5 | - | - | - | 5 (100%), 0 (0%) | - |
| **Combined Total**<br>**(n = non-missing**<br>**value count)** |  | **2,251** | **63.0 [52.0, 73.0]**<br>**(n = 1,807)** | **1,145 (58%)**<br>**(n = 1,969)** | **Black: 120 (36%)**<br>**Asian: 61 (18%)**<br>**White: 132 (39%)**<br>**Other: 22 (7%)**<br>**(n = 335)** | **1,887 (84%), 364 (16%)**<br>**(n = 2,251)** | **1,307 (76%)**<br>**(n = 1,710)** |

### Molecular Subtyping

Molecular subtypes were defined using cluster analysis of 5,000 genes, including immune-related genes, lipid genes, and the most variable genes in the dataset (Materials and Methods Supplement). This approach was used to guide molecular subtype identification based on known immune-related pathways/processes, because most gene expression variability (>90%) in the dataset is captured within these genes (Supplemental Figure 1), and to minimize confounding factors such as sex, age, and genetic ancestry.

We compared performance of three clustering methods using ConsensusClusterPlus (Version 1.64)^34^: K-means, partition around medoids (PAM), and hierarchical clustering. Each method was repeated 1,000 times with a resampling proportion of 0.8 and a range of clusters from 2-7. All clustering approaches showed K=4 as the optimal number of clusters (Supplemental Table 2). We ultimately selected the K-Means clustering approach to derive the sepsis molecular subtypes due to its greater silhouette widths and higher cluster consensus (Material and Methods Supplement, Supplemental Table 2).

### Enrichment in the Molecular Subtypes

To identify molecular subtype-specific genes and biological pathways, we performed pairwise differential expression analysis between all pairs of molecular subtypes. A gene was assigned to a molecular subtype if it was significantly differentially expressed in that molecular subtype in the same direction (up- or down-regulated) in each comparison (Supplemental Table 3). Genes assigned to each molecular subtype were then used to identify molecular subtype-associated pathways using Biological Process ontologies with the global network in HumanBase. Up- and down-regulated genes for each molecular subtype were queried separately and only pathways with Q-values <= 0.05 were retained (Supplemental Table 4).

### Identifying Molecular subtype-specific Drug Repurposing

We next performed enrichment analysis of drug-gene relationships in molecular subtype signatures. To do this, we extracted drug-gene relationships from the ClinPGx pharmacogenomic database (downloaded on June, 16^th^ 2025) for the top 50 differentially expressed genes in each molecular subtype (Table 4, Supplemental Table 6).

### Analysis of Mortality Data

Given that time-to-event survival data and mortality were not included in all datasets, we analyzed available mortality data in three ways. First, using mortality as a binary variable for all patients with which we had mortality data (n = 1,710), we performed differential expression analysis between survivors and non-survivors using Limma (v 3.56.2)^35^, followed by multiple hypothesis correction on differential expression results (Figure 4A,C). Out of the 5,000 genes we use for analysis, 1,825 genes were differentially expressed between survivors and non-survivors (Supplemental Table 3). Notably, all HLA genes in our dataset were downregulated in non-survivors.

We next performed restricted mean survival time (RMST) analysis of 28-day mortality for all patients with which we had complete time-to-event data (560 patients) using the survival package(v 3.8)^36^ and ggsurvfit (v1.1.0)^37^ packages. For each molecular subtype, RMST was estimated in a two-group comparison by comparing the time-to-event status at 28-days of one molecular subtype to the average of time-to-event status at 28-days of the other molecule subtypes (Figure 4B). Statistics were calculated based on these results. Only C4 had a significant difference in RMST after BH correction, with patients in C4 having worse survival.

In our final mortality analysis, using mortality as a binary variable, we compared mortality status in samples with mortality data (n = 1,710) across our four molecular subtypes. To do this, we used a two-way table of mortality status after 28 days and performed association tests using contingency table analysis using survRM2 (v 1.0)^38^. We found statistically significant associations between mortality status and our four molecular subtypes (BH-corrected Chi-squared test p-value = 1.4e-07). This significance was primarily driven by the decrease in mortality of patients in C2, with only 9% of non-survivors being assigned to C2 (Figure 3D). We also found higher than expected mortality in C1 and C4, with worse 28-day survival outcomes (Figure 4B).

**Figure 3.**
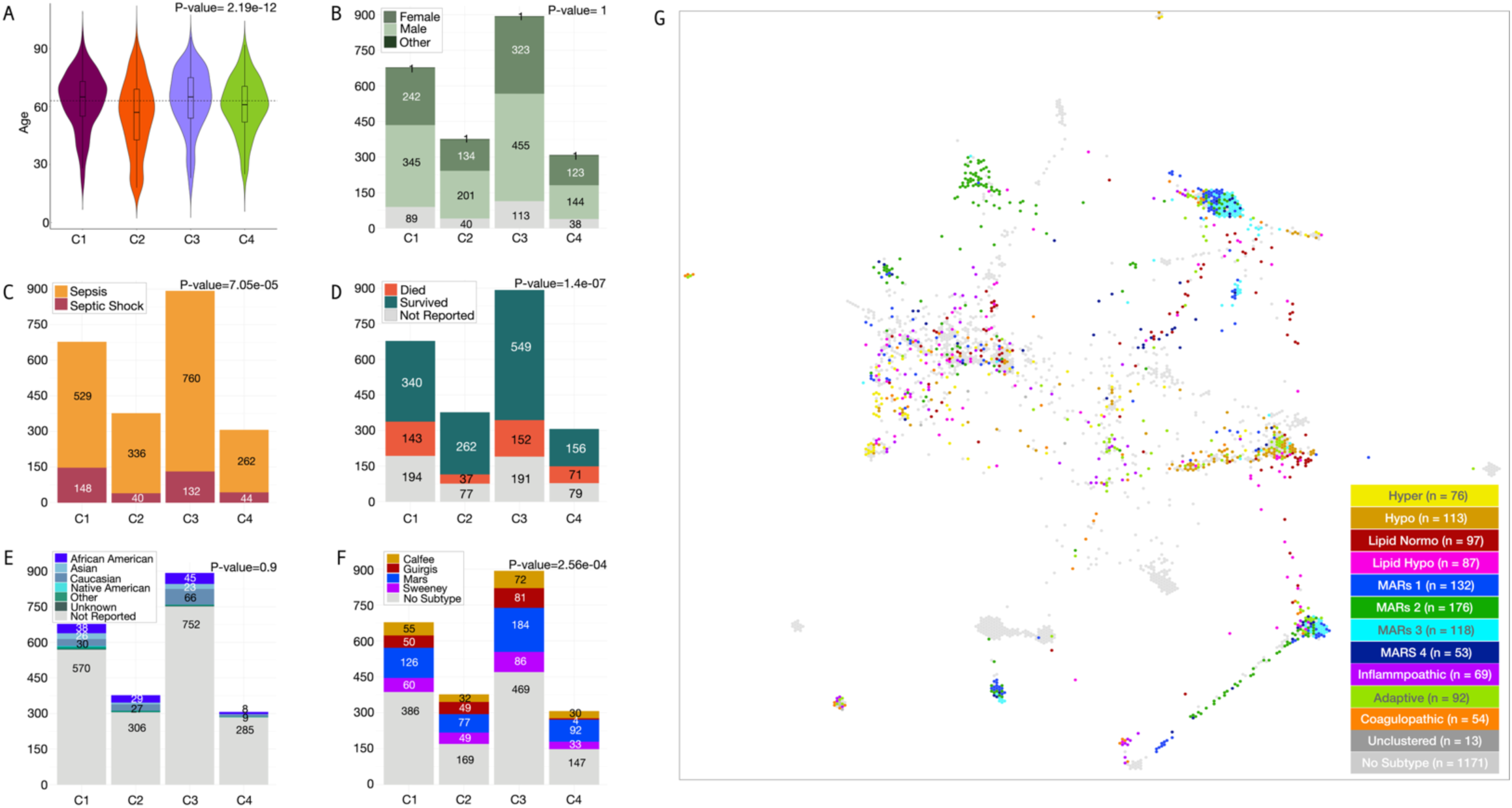
Molecular subtype Clinical Enrichment. (A) Violin plots with box plots showing age distribution of each molecular subtype. The dotted horizontal line marks the median age across all molecular subtypes. (B-F) Barplots for clinical variables used in analysis showing distribution of each variable across molecular subtypes. (G) TumorMap showing samples colored by previously defined molecular subtype (if applicable). A TumorMap visualizes pairwise sample-sample molecular similarity within a group of samples across a hexagonal grid in a force-directed and non-overlapping manner and provides a unique overview of disease structure that has not been previously applied to sepsis.

**Figure 4.**
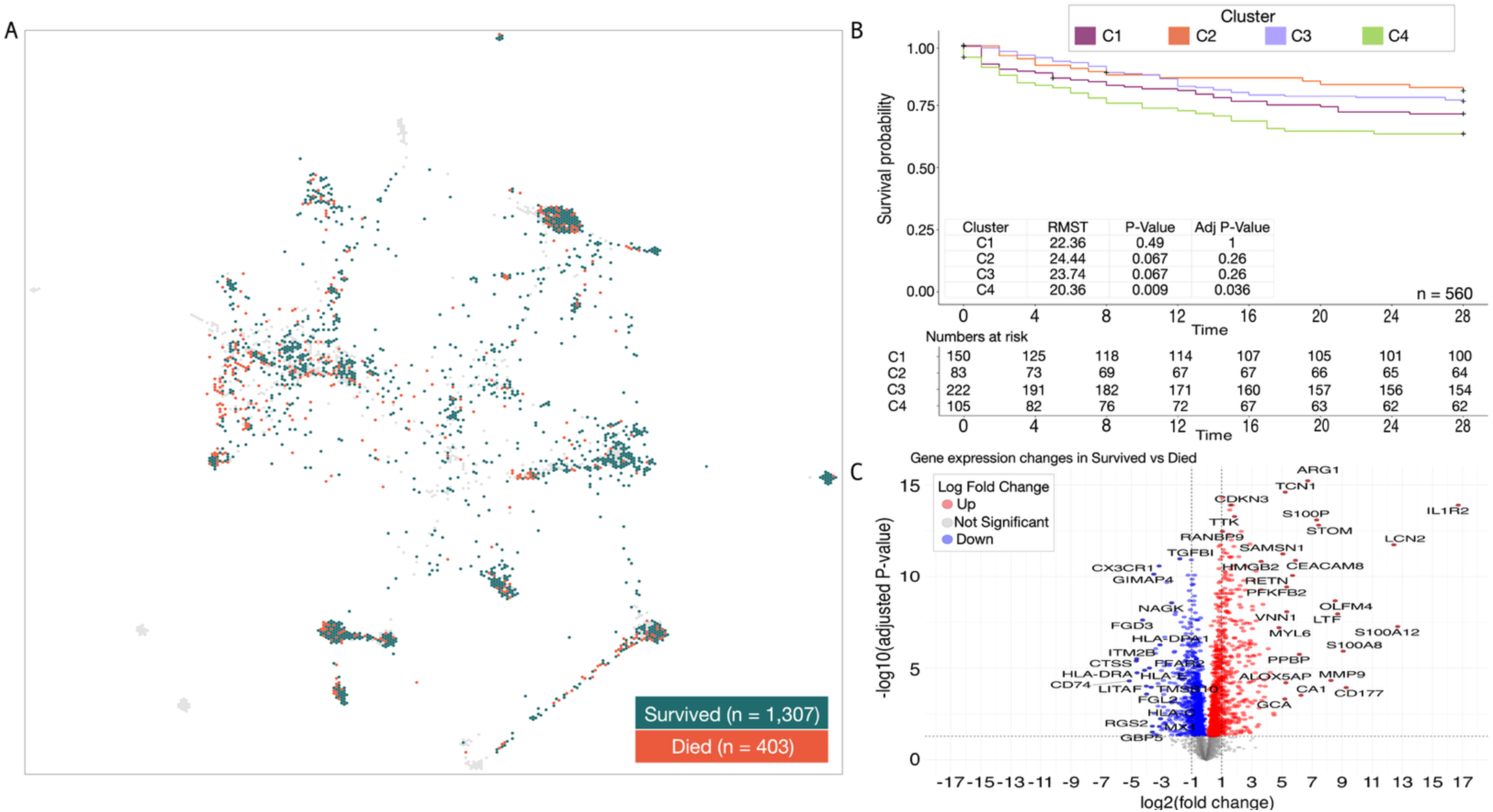
Analysis of Mortality Data. (A) TumorMap of all samples colored by mortality in patients with available mortality data (n = 1,710). The TumorMap is the same as depicted in Figure 3G but colored by survival. (B) Kaplan Meier survival plot showing survival probability by molecular subtype for patients with available time to mortality data (n = 560). RMST p-values show difference between molecular subtype time to survival VS all other molecular subtype time to survivals. (C) Volcano plot of differentially expressed genes between survival and mortality for patients with available mortality data (n = 1,710).

## Results

### Deciphering sepsis molecular subtypes

Cluster analysis identified four molecular subtypes (C1 = 677, C2 = 376, C3 = 892, C4 = 306 samples) from the sepsis data atlas (see Methods). The molecular subtypes had overall low cluster cohesiveness metrics, which is concordant with previous studies and expected given the heterogeneity of sepsis^39,40^. However, molecular subtypes remained stable across multiple clustering iterations when using our approach of biologically-guided gene selection to minimize molecular confounding factors, which improved molecular subtype robustness. This is demonstrated visually in the Figure 2A t-SNE plot.

### Molecular subtype-specific gene set enrichment analysis

Gene signatures for each molecular subtype were defined based on pairwise expression comparisons of all molecular subtypes (see Methods). This resulted in 301-1,067 genes per molecular subtype signature. Gene set enrichment analysis identified 820 unique pathways for C1, 627 unique pathways for C2, 177 unique pathways for C3, and 637 unique pathways for C4. Overall, gene set enrichment analysis results were primarily defined by their uniqueness in pathways associated with MHC class II functions, DNA damage, homeostatic pathways, and coagulation (see Supplemental Table 4). Figure 2B shows a heatmap of the top 25 most differentially expressed genes for each molecular subtype and highlights unique expression patterns across molecular subtypes. Full results are included in Supplemental Table 4 and summarized in the following paragraphs.

C1 showed marked by downregulation of both innate and adaptive immune response pathways, including interferon signaling, T and natural killer (NK) cell activation, and antigen presentation. C1 has increased expression of proinflammatory cytokines (notably IL-1, IL-6). Lipid metabolism in C1 was dysregulated, with upregulated pathways related to lipid storage and ATP generation; in addition to increased reactive oxygen species and proteasomal protein catabolism. Overall, the C1 molecular subtype was characterized by immune exhaustion, impaired pathogen clearance, and metabolic dysregulation.

C2 had the lowest mortality rate of all molecular subtypes. C2 displayed broad upregulation of immune pathways, including upregulation of IL-2, IL-4, IL-10, IL-13, IL-17, and interferons, supportive of robust, adaptive, and efficient immune response. Additionally, pathways indicated increased activation and differentiation of T and B cells, upregulation of pathways related to metabolic maintenance, enhanced antigen processing and presentation, cell signaling, and increased expression of genes related to telomere protection demonstrate a balanced immune response. C2 showed a strong, coordinated immune response and metabolic adaptability, which may partly explain improved survival compared to other molecular subtypes.

C3 was defined primarily by the upregulation of pathways maintaining cellular homeostasis (oxygen homeostasis, angiogenesis) and response to cellular and oxidative stress. C3 had elevated levels of proinflammatory factors such as IL-1, IL-6, response to lipopolysaccharide (LPS), activation of JAK/STAT and MAPK pathways, increased chemokine production, and G-protein coupled receptor (GPCR) signaling. C3 was also characterized by heightened cellular stress responses and controlled inflammation, which may support tissue repair but risk organ dysfunction.

C4 had the highest mortality of our molecular subtypes. C4 was predominantly defined by suppression of innate and adaptive immune responses, broad dysregulation of lipid and carbohydrate metabolism, and upregulation of pathways related to protein degradation. Additionally, C4 had increased coagulation, cellular apoptosis, and oxidative stress responses. Immune signaling pathways such as cytokines IL-1β, IL-6, IL-8, IL-10, IL-12, IFN-γ and toll-like receptor signaling were downregulated in C4. Ultimately, C4 represented an immunosuppressed molecular subtype with metabolic reprogramming and coagulopathy indicative of higher mortality.

### Molecular Subtype Demographic Enrichment

To identify demographically relevant information for each molecular subtype, we statistically tested correlations between our molecular subtypes and demographic data that were available for at least 20% of the samples (Supplemental Table 5). All 2,251 samples had a sepsis or septic shock diagnosis (1,887 sepsis, 364 septic shock). We found a statistically significant association between proportions of sepsis/septic shock within each molecular subtype (Benjamini-Hochberg (BH) corrected Chi-square test p-value = 7.05e-05), which was primarily driven by a higher concentration of septic shock patients in C1 and a lower concentration of septic shock patients in C2 (Figure 3C). Age was available for 1,807 samples from 15 of the 28 studies and was significantly associated with our molecular subtypes (BH corrected one-way ANOVA, p-value = 2.19e-12), with the strongest association found in C2, which had the youngest patient population and highest survival (Figure 3A). Biological sex classification was available from 18 of the studies, totaling 1,967 samples. Biological sex was not significantly associated with molecular subtype (BH corrected Chi-squared test p-value = 1, Figure 3B). Race data was available for 335 samples in 3 datasets and was not statistically significantly associated with our molecular subtypes (BH corrected Fisher’s exact test p-value = 0.9, Figure 3E).

Next, we tested associations between our molecular subtypes and four sets of previously published sepsis subtypes (Figures 3F-G and Supplemental Figure 2)^31,39,40,42–44^. Of the 2,251 sepsis samples in the data atlas, 1,080 (48%) had a previously assigned subtype from one of these studies. We found statistically significant enrichment across all previously defined subtypes within our molecular subtypes (Table 3, Supplemental Figure 2), highlighting how previous studies identified similar patterns in sepsis heterogeneity but that the larger data atlas presented here was essential to linking together these molecular patterns in sepsis patients.

**Table 3.** Comparison to Previously Defined Subtypes. Results from contingency table analysis of our four identified molecular subtypes with previously defined subtypes.

| Study | Included Samples | Subtypes | BH-Corrected P-value | Molecular Subtype association (~) |
| --- | --- | --- | --- | --- |
| Scicluna et al.<sup>39</sup> | 479 | Mars1, Mars2, Mars3, Mars4 | 1.34e-89 | Mars1 ~ C4, Mars2 ~ C1, Mars3 ~ C3 and C2 |
| Sweeney et al.<sup>40</sup> | 327 | Inflammopathic, adaptive, coagulopathic | 4.99e-03 | Adaptive ~ C2 and C3, Coagulopathic ~ C3, Inflammopathic ~ C1 |
| Calfee et al., Neyton et al.<sup>42,44</sup> | 189 | Hyperinflammatory, hypoinflammatory | 8.77e-07 | Hyperinflammatory ~ C1, Hypoinflammatory ~ C3 |
| Guirgis et al., Augustin et al.<sup>31,43</sup> | 184 | Hypo_L, Normo_L | 3.78e-03 | Normo-L ~ C3, Hypo-L ~ C1 |

**Table 4.**
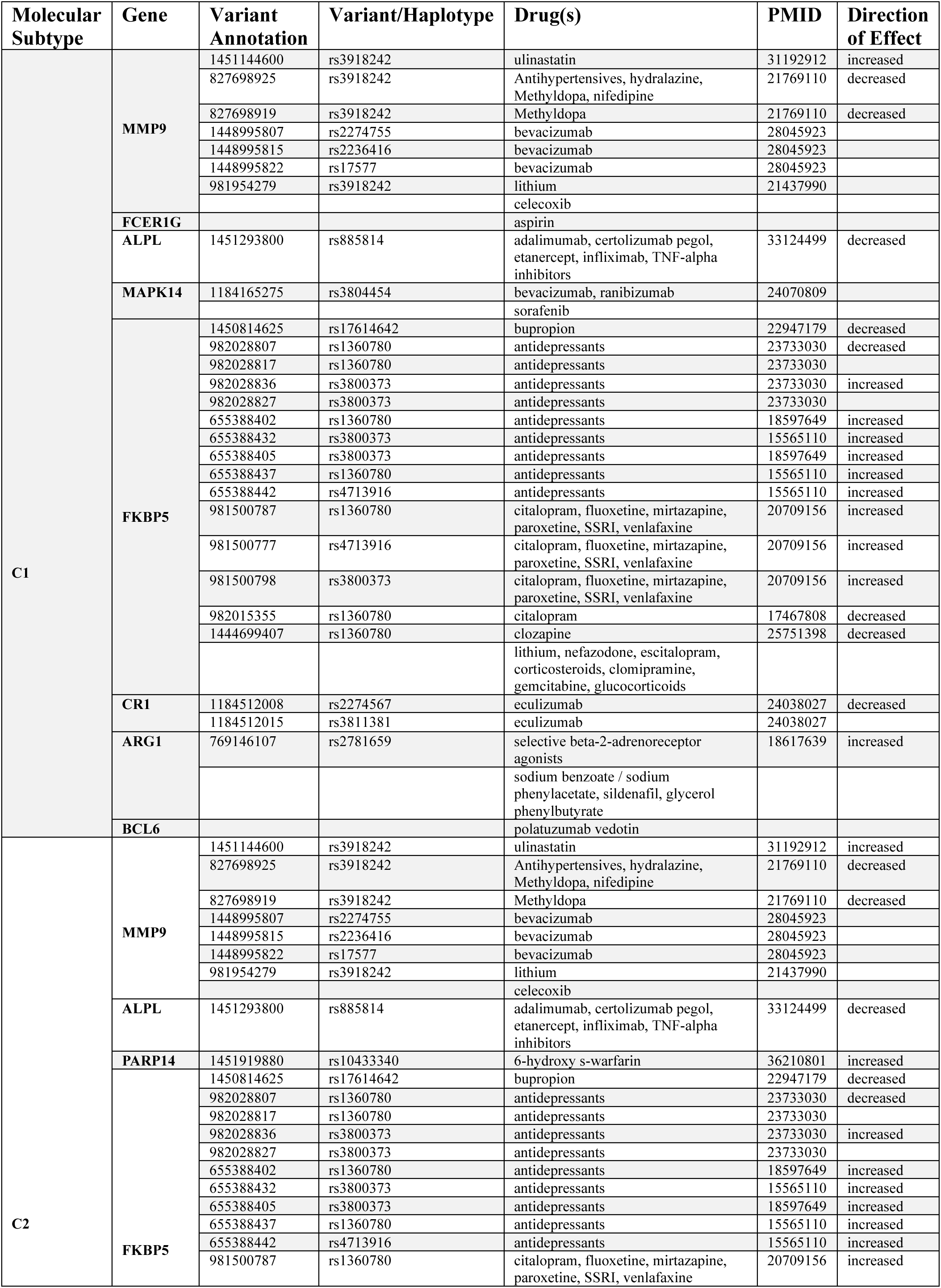
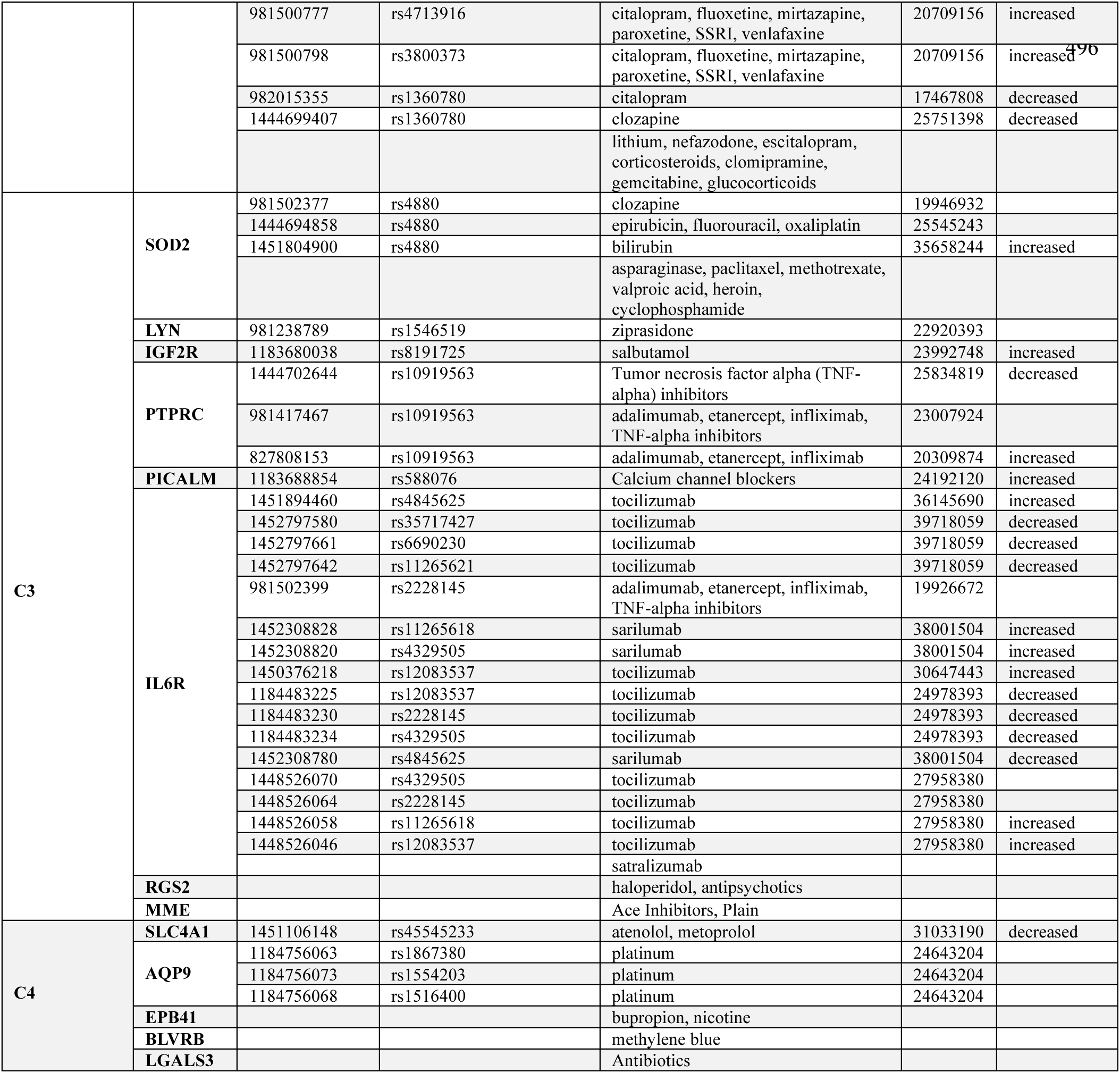
ClinPGx Variant, Gene and Drug Relationship Data. Variant, gene and drug relationship data for each molecular subtype downloaded from ClinPGx.

### Analyzing Mortality Data

Differential expression analysis between survivors and non-survivors (Figure 4A,C) identified 1,825 differentially expressed genes (Supplemental Table 3). All HLA genes in our dataset were downregulated in non-survivors. Pathway enrichment analysis of differentially expressed genes identified 1,234 unique pathways (Supplemental Table 4). Restricted mean survival time (RMST) analysis of 28-day mortality for all patients with complete time-to-event data revealed significant differences in RMST for C4 (Figure 4B). Contingency table analysis between survivors and non-survivors and our molecular subtypes showed statistically significant associations between mortality status and our four molecular subtypes (BH-corrected Chi-squared test p-value = 1.4e-07), primarily driven by the relatively decreased mortality observed in C2 (Figure 3D). We also found higher than expected mortality in C1 and C4, with worse 28-day survival outcomes (Figure 4B). Additional details regarding mortality data analysis can be found in the Material and Methods Supplement.

### Identifying Potential Drugs for Each Molecular Subtype

Molecular sepsis subtypes may help explain failed clinical trials and advance precision medicine. If subtypes are not considered during treatment assignment, the assignment of patients who are unlikely to respond to the treatment being investigated can confound clinical trial results. To address this and to identify potential new treatment by sepsis molecular subtype, we searched the ClinPGx database for the top 50 differentially expressed genes from each molecular subtype (Table 4 and Supplemental Table 6). This resulted in 144 unique genes, 22 of which had existing drugs that modulate them. Analysis of these gene–drug associations across our molecular subtypes revealed distinct therapeutic possibilities by molecular subtype (Table 4).

C1 was characterized by enrichment of genes such as MMP9, FCER1G, MAPK14, and CR1, corresponding drug associations included anti-inflammatory and immunomodulatory agents such as aspirin, celecoxib, sorafenib, and eculizumab, as well as broader classes like corticosteroids, antidepressants, and adrenergic modulators. C2 showed overlap with MMP9 and FKBP5, reinforcing links to anti-inflammatory drugs (celecoxib) and psychiatric medications (e.g., SSRIs, SNRIs, lithium), suggesting shared inflammatory and stress-response pathways. In C3, gene associations included RGS2, SOD2, MME, PTPRC, and PICALM, with therapeutic connections spanning diverse domains: antipsychotics (haloperidol), cytotoxic chemotherapies (asparaginase, paclitaxel, methotrexate), cardiovascular drugs (ACE inhibitors, calcium channel blockers), and immunomodulators (infliximab, etanercept, tocilizumab), highlighting the interplay between immune regulation, oxidative stress, and cardiovascular signaling. Finally, C4 was linked to SLC4A1, EPB41, BLVRB, and LGALS3, genes associated with drug classes such as beta-blockers (atenolol, metoprolol), antidepressants (bupropion), redox-modulating agents (methylene blue), and antibiotics, indicating a distinct molecular profile related to erythrocyte stability, oxidative stress, and microbial defense.

## Discussion

Despite progress in clinical care, treatment for sepsis remains challenging due to the high degree of syndrome heterogeneity. Large-scale genetic studies are a promising approach for uncovering the molecular mechanisms underlying sepsis and guiding the development of targeted therapies. Here, we created a large-scale transcriptomic atlas of 28 sepsis studies, from which we identified 4 distinct sepsis molecular subtypes that may provide guidance for precision medicine in sepsis (Table 5).

**Table 5.** Profiling of Molecular Subtypes. Key characteristics of identified molecular subtypes.

| Molecular Subtype | Key Biology & Risk | Potential Clinical Actions | Candidate Drugs / Interventions |
| --- | --- | --- | --- |
| C1 – Immune exhaustion + metabolic dysregulation (↑ mortality, shock) | ↓ MHC II, T/NK dysfunction<br>↑ IL-1/IL-6 (cytokine storm)<br>Dyslipidemia, ARG1 ↑, MMP9 ↑ | Early ICU admission<br>Monitor for shock | Corticosteroids<br>Anti-MMP9 (bevacizumab)<br>Eculizumab (CR1)<br>ARG1 modulators<br>Aspirin, celecoxib, sorafenib |
| C2 – Robust adaptive immunity (lowest mortality, younger patients) | ↑ IL-2, IL-4, IL-10, IFN<br>Balanced pro/anti-inflammation<br>Telomere protection, JAK/STAT ↑ | Standard care<br>Supportive therapy | Celecoxib, SSRIs/SNRIs, lithium<br>Bevacizumab (MAPK14) |
| C3 – Inflammatory & cellular stress (intermediate mortality) | ↑ JAK/STAT, MAPK, LPS response<br>↑ IL-1, IL-6, chemokines<br>Lysosome & platelet activation ↑ | Monitor for organ dysfunction<br>Platelet/coagulation surveillance | Anti-IL6R biologics (tocilizumab, sarilumab, infliximab, adalimumab)<br>GM-CSF |
| C4 – Immunosuppressed, metabolic reprogramming (highest mortality) | ↓ innate/adaptive immunity<br>Dyslipidemia, glycolipid/sphingolipid ↑<br>↑ Coagulation, apoptosis, protein degradation | High-risk ICU triage<br>Monitor refractory shock & coagulopathy<br>Consider vasopressor adjuncts | Methylene blue (BLVRB/redox)<br>Antibiotics |

Of all molecular subtypes, C2 had the youngest patient population and best survival. Pathways activated in C2 were reflective of an effective and protective immune response. In contrast, C4 showed widespread immune suppression and was associated with the highest mortality. Patients with this molecular subtype will likely benefit the most from early ICU admission and intervention. Following C4, C1 was associated with the second highest mortality and demonstrated severe immune dysregulation with a greater proportion of septic shock patients than expected. C3’s pathways were suggestive of a responsive immune state with preserved regulation and an intermediate mortality. An overview of molecular subtype characteristics, drug-relationships and clinical implications is provided in Table 5.

Our molecular subtypes correlated with previously defined subtypes, with parallels in sample-subtype assignment, gene and pathway enrichment, and survival trend. Mars1-4 subtypes, defined by Scicluna et al.^39^ based on unsupervised clustering of transcriptomic data, with Mars1 having the highest mortality, followed by Mars4, Mars3, and Mars2 (lowest mortality). Another study by Sweeney et al.^40^ defined three subtypes (Inflammopathic, Adaptive and Coagulopathic) based on unsupervised clustering of 14 previously published transcriptomic datasets, with Coagulopathic and Inflammopathic having higher patient mortality. Calfee et al.^42^ defined Hyperinflammatory (higher mortality) and Hypoinflammatory subtypes based on clinical and plasma protein biomarker data in ARDs patients and further applied to transcriptomic data by Neyton et al.^44^ Guirgis et al.^31^ identified two subtypes Hypolipoprotein (Hypo-L) and Normolipoprotein (Normo-L) based on patient lipoprotein levels, with Hypo-L subtype having higher mortality. Our C1 molecular subtype correlated with Mars2, Inflammopathic, Hyperinflammatory, and Hypo-L subtypes from previous studies, with shared enrichment in pathways related to inflammatory cytokines, cell recognition, lipid dysregulation and NF-κB (Supplemental Figure 2) ^31,39,40,42,44^. C2 had the highest survival and correlated with previous subtypes enriched for immune signaling pathways such as Mars3, Mars4, and Adaptive subtypes. C3 correlated with Mars3, Hypoinflammatory, and Normo-L subtypes with less inflammation and increased cellular response and homeostasis. C4 correlated with Mars1 and Coagulopathic subtypes, which had the highest mortality rates out of their respective subtypes and had similar downregulation in Toll-like receptor, NF-κB, and T-cell signaling. Across comparisons we observed that higher mortality molecular subtypes had less adaptive immunity and more inflammation.

Our molecular subtypes closely correlated with 28-day mortality. This significance was driven by the relatively lower mortality of patients in C2 (Figure 3D) compared to the increased mortality of patients in C1 and C4 (Fig4 B). Comparison of differentially expressed genes and enriched pathways across molecular subtypes and survival status revealed that C1 exhibited the greatest similarity in pathway regulation to non-survivors, followed by C4. C2 had the most similar gene expression to survivors with 38% of genes upregulated in patients who survived being similarly upregulated in C2. These genes included CTSS and CD74, both of which have previously been shown to be promising biomarkers in sepsis prognosis (Figure 4C)^45,46^. Similarly, 12% of down-regulated genes in C2 were downregulated in survivors, including the most significantly differentially expressed genes IL1R2, LCN2, and STOM (Figure 4C & Figure 5B). Sixty-eight percent of C2 pathways were similarly regulated in survivors, such as upregulation of proinflammatory interleukins and downregulation of TGFβ receptor signaling.

**Figure 5.**
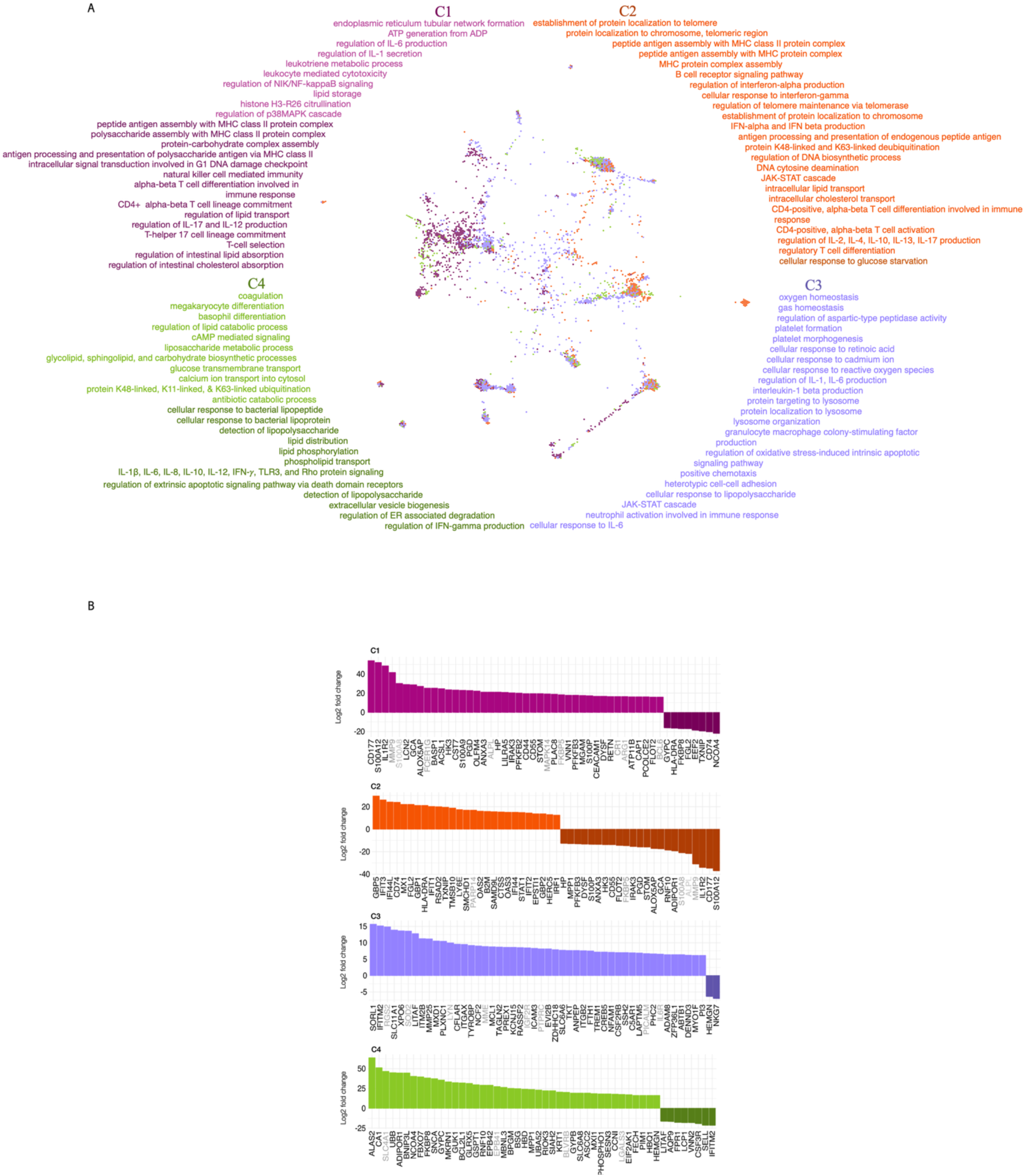
Pathway enrichment and drug associations for sepsis molecular subtypes. (A) TumorMap colored by our four defined molecular subtypes with associated up-regulated and down-regulated pathways for each molecular subtype. The TumorMap is the same as depicted in Figure 3G but colored by our defined molecular subtypes. For each molecular subtype’s pathways, lighter colors indicate upregulation and darker colors indicate down regulation. (B) Barplots for the top 50 most differentially expressed genes for each molecular subtype in comparison to the average expression of the other three molecular subtype.

Differential expression analysis of sepsis versus septic shock patients revealed several genes that repeatedly appear to be enriched among poorer outcome patients. For example, genes CD177, S100A12, IL1R2, MMP9, LCN2 were consistently among the most differentially expressed genes, being upregulated in higher shock and mortality clusters (C1or C4, septic shock, mortality). One particularly relevant gene, Arginase-1 (ARG1) has emerged as a promising biomarker for both disease severity and treatment response in conditions such as sepsis and septic acute respiratory distress syndrome (ARDS) patients^47^. Arginases have been implicated in many diseases and are involved in dysfunction across organ systems. In the immune system, overexpression of ARG1 inhibits T-cell activity and promotes immune suppression, leading to poorer clinical outcomes. This suggests that for certain sepsis patients, targeted modulation of ARG1 may be an effective clinical management approach.

For drug repurposing, there was, in general, a significant representation of monoclonal antibody therapies across molecular subtypes. These types of drugs are gaining more attention in fighting bacterial pathogens, especially in sepsis, where heterogeneity in disease progression and host response necessitates precision targeting^48^. In C1, targeting MMP9 via acizacizumab is particularly promising and worthy of further investigation. MMP9 is overexpressed in C1, non-survivors, and septic shock, suggesting a negative effect on patient survival. Previous studies have found conflicting results regarding the roles of MMP9 and its inhibitor TIMP-1 in sepsis^49^, however due to its consistent upregulation in C1 and septic shock and non-survivors in this study it warrants further investigation. Bevacizumab was a target for C2 via MAPK14, which was differentially expressed in both C1 and C2. Bevacizumab is primarily used for treatment in cancers in combination with other drugs^50,51^. It is important to note that genes in C1 and C2 are often inversely correlated, therefore a drug used to target gene expression in one subtype will likely not be an effective treatment in the other. This has important implications for the design of future clinical trials, and this concept may explain why prior trials with broad inclusion criteria have failed. Additionally, Eculizumab may be a promising target as it decreases expression of CR1 in C1, which plays an important role in the innate immune system and has been shown to be overexpressed in septic patients vs controls^52^. Case reports of sepsis treatment with eculizumab show it may have promise in certain sepsis cases, but must be closely monitored^53,54^. Adalimumab (ALPL, PTPRC, IL6R) and infliximab (PTPRC, ALPL, IL6R), are targets in C1, C2, and C3. Infliximab has been identified as having a positive response in a subtype of patients with ulcerative colitis (UC) and Crohn’s disease (CD), diseases related to immune dysfunction^16,30^. Bevacizumab, infliximab, and adalimumab have gene targets in differentially expressed genes in survivor vs non-survivor and sepsis vs septic shock comparisons. Tocilizumab and sarilumab modulate IL6R in C3 and have undergone some investigation in the treatment of sepsis and COVID-19 with mixed results^55–58^. These drugs are IL-6 receptor antagonists, which allow them to block IL-6 binding and inhibit downstream immune and inflammatory responses. Tocilizumab has also been of interest in treating pediatric sepsis/septic shock^59^.

In addition to monoclonal antibody drug targets, C1 is a potential target for corticosteroids via modulation of FKBP5 as well as glycerol phenylbutyrate and sodium benzoate / sodium phenylacetate via ARG1. Corticosteroids have previously been shown to have differential benefit depending on transcriptomic profile at the time of septic shock onset^17^. Subtypes characterized by T cell exhaustion and downregulation of MHC II antigens, both features of C1, are shown to have increased benefit from steroids. Additionally, ARG1 has been identified as a promising biomarker and therapeutic target in sepsis and there is an ongoing investigation in the repurposing of FDA-approved drugs for modulation of ARG1^60^. C4 had upregulation of BLVRB, a gene which plays an important role in redox-regulated mechanisms. These mechanisms affect hematopoietic stem cells, platelets, and, in turn, blood clotting, distinct characteristics of C4. Methylene blue acts on BLVRB and has been shown to decrease patient stays in the ICU, reduce days on mechanical ventilation, reduce time on vasopressors, and increase vasopressor-free days^61^. Methylene blue is sometimes used to treat patients with refractory septic shock in the ICU, though clear guidance on its use has not been established.

### Study Limitations

While our study objective was to derive transcriptomic molecular subtypes within sepsis, it was limited by the availability of clinical variables across studies. Due to this, we had limited ability to investigate clinical differences between molecular subtypes. Our study was also limited due to the potentially different definitions and inclusion criteria for sepsis in each included study, however we addressed this by reviewing inclusion criteria used for each study as well as study specific diagnoses, excluding samples with known parasitic infections or other unique cases of sepsis, such as melioidosis.

### Conclusions

The rapid progression and inherent heterogeneity of sepsis make the development of effective treatments challenging. Integration of transcriptomic data into disease modeling has been widely successful in identifying molecular features predictive of patient prognosis and treatment response. Studies of gene expression data in sepsis have been limited to relatively small samples sizes, which have limited the development of reproducible subtypes across patient cohorts and subsequent treatment development.

Here we provide a comprehensive transcriptomic analysis of sepsis through a meta-analysis of adult sepsis patients. We created a sepsis data atlas consisting of 2,251 unique sepsis/septic shock patients, evaluated patterns of gene expression and identified molecular subtypes. Using this atlas, we uncovered 4 molecular subtypes (C1-C4), described their distinct expression patterns and molecular pathways, and identified potential biomarkers and therapeutic targets relevant to each. Our detailed molecular analysis identifies potential drug targets within each molecular subtype with implications for future precision medicine for sepsis. The ability to accurately identify the clinically-relevant molecular subtypes presented here would provide a potential avenue for improving sepsis outcomes and clinical management.

## Author contributions

Conceptualization: Kiley Graim and Faheem Guirgis

Methodology: Kiley Graim and Leslie A. Smith

Investigation: Kiley Graim, Faheem Guirgis, Lauren P. Black and Leslie A. Smith

Visualization: Kiley Graim and Leslie A. Smith

Supervision: Kiley Graim and Faheem Guirgis

Writing—original draft: Kiley Graim, Faheem Guirgis, Srinivasa T. Reddy, Leslie A. Smith Writing—review: Leslie A. Smith, Beulah Augustin, Vinitha Jacob, Lauren P. Black, Andrew

Bertrand, Charlotte Hopson, Emilio Cagmat, Susmita Datta, Srinivasa T. Reddy, Faheem W. Guirgis, Kiley Graim

Writing —editing: Leslie A. Smith, Beulah Augustin, Vinitha Jacob, Lauren P. Black, Andrew Bertrand, Charlotte Hopson, Emilio Cagmat, Susmita Datta, Srinivasa T. Reddy, Faheem W. Guirgis, Kiley Graim

## Impact Statement

To our knowledge, this work represents largest combined sepsis transcriptomic dataset to date and identifies robust molecular subtypes in sepsis. It provides an in-depth gene expression and pathway analysis, identifying characteristics that may discriminate sepsis patient subpopulations at the transcriptomic level.

## Acknowledgments

This work was supported by funding from NIH/NIGMS R01GM133815, NIH/NIAID R21AI188477, NIH/NIGMS K08 K08GM151392 (V.J.), and a 2023 UF AI2Heal Datathon Award. LAS is supported by a UF Dean’s Fellowship Award and by the UF GenNext program. This project makes use of publicly available data from GSE65682, GSE185263, GSE236892, GSE134347, GSE236713, GSE189400, GSE131761, GSE74224, GSE66890, GSE95233, GSE32707, GSE154918, GSE216902, GSE63311, GSE57065, GSE13015, GSE137340, SRP198776, GSE232753, GSE33118, GSE69063, GSE196117, GSE199816, GSE100159, GSE211210, GSE222393, GSE232404, and Guirgis et al^31^. This manuscript is the result of funding in whole or in part by the National Institutes of Health (NIH). It is subject to the NIH Public Access Policy. Through acceptance of this federal funding, NIH has been given a right to make this manuscript publicly available in PubMed Central upon the Official Date of Publication, as defined by the NIH.

## Data and materials availability

Gene expression matrix and metadata for all samples in all included studies provided on GEO under accession GSE310929. Data processing scripts provided on GitHub at github.com/leslie-smith1112/SepsisPhenotyping/tree/main.

## Supplemental Tables

Complete Supplemental Tables are located in Sepsis_Final

**Supplemental Table 1. Harmonized metadata for all samples included in the data atlas.** Harmonized metadata from all datasets included in the data atlas. This harmonized dataset includes all samples (3,713 samples) used for initial batch correction.

**Supplemental Table 2. Cluster analysis method comparison.** Results from the clustering runs of the three clustering methods compared in this analysis (K-Means, PAM, Hierarchical Clustering).

**Supplemental Table 3. Gene signatures for each molecular subtype.** Differential expression results for the genes assigned to each molecular subtype. Genes assigned to each molecular subtype were similarly differentially expressed in all pairwise comparisons expression comparisons (see Methods).

**Supplemental Table 4. Molecular subtype pathway enrichment.** Pathway enrichment results for each molecular subtype. Pathway analysis was done using Biological Process terms in HumanBase using the global network.

**Supplemental Table 5. Statistical Analysis.** All results from statistical analyses done on identified molecular subtypes. This includes contingency table analysis of available demographic data as well as comparisons to previously identified subtypes. For each test done on a subset of samples with available data, a null distribution is provided.

**Supplemental Table 6. Molecular subtype-specific drug-gene relationships.** The top 50 most differentially expressed genes in each molecular subtype were queried in ClinPGx to identify possible molecular subtype-specific drug associations. A smaller version of this table is provided in the main manuscript (Table 4), this provides all available information from the drug queries.

**Supplemental Table 7. Included dataset summaries and links.** List of all studies included in the data atlas with brief summaries of the study, sequencing type, date published, and paper links.

## Supplemental Figure Legends

**Supplemental Figure 1:**
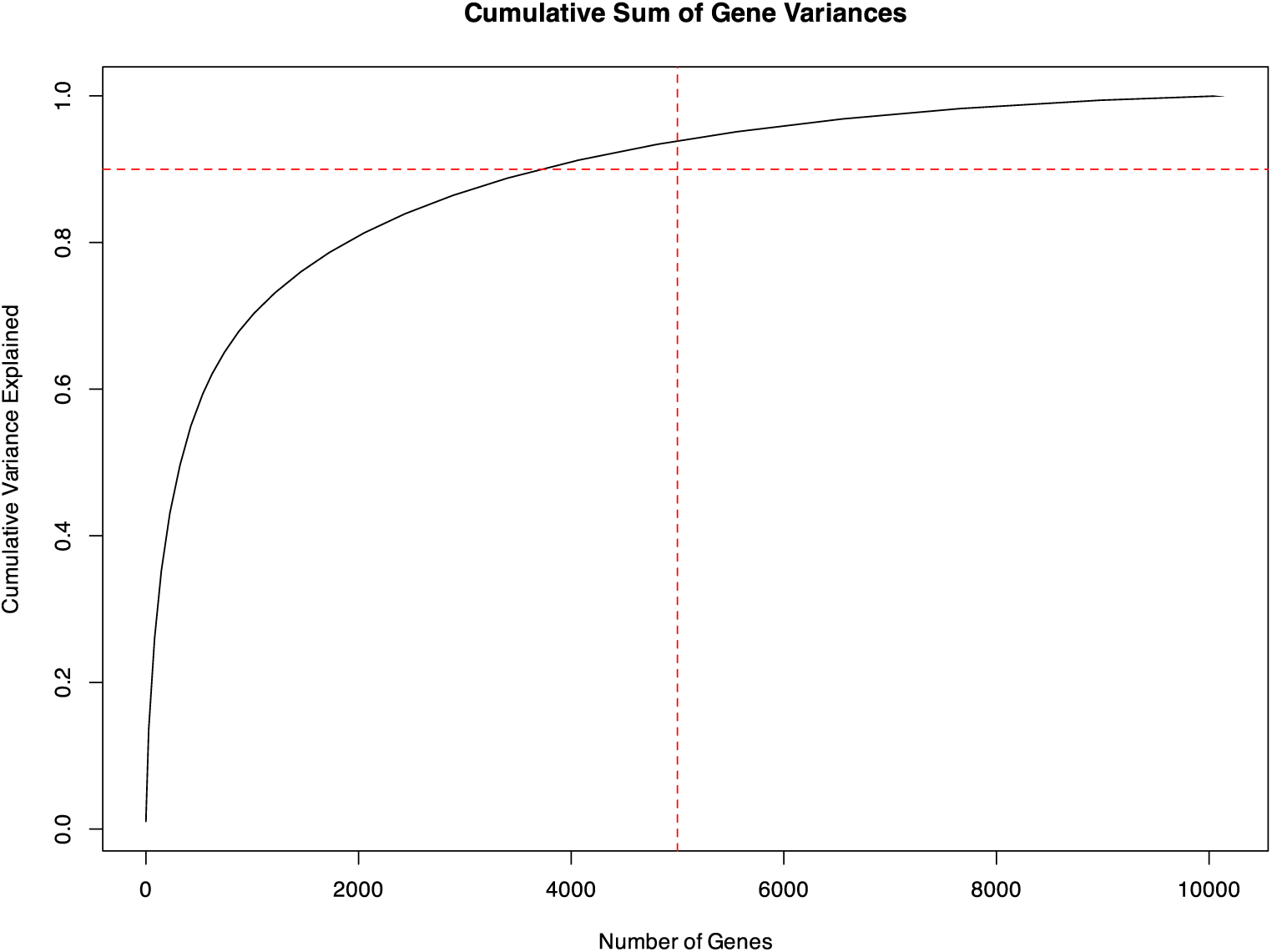
Cumulative dataset variance. The relative percent of variance in the dataset captured within a subset of genes. A horizontal line shows where 90% of variance is captured and a vertical line shows 5,000 gene cut off used in the manuscript. Using 5,000 genes captures more than 90% of the total variability in the transcriptomic data.

**Supplemental Figure 2:**
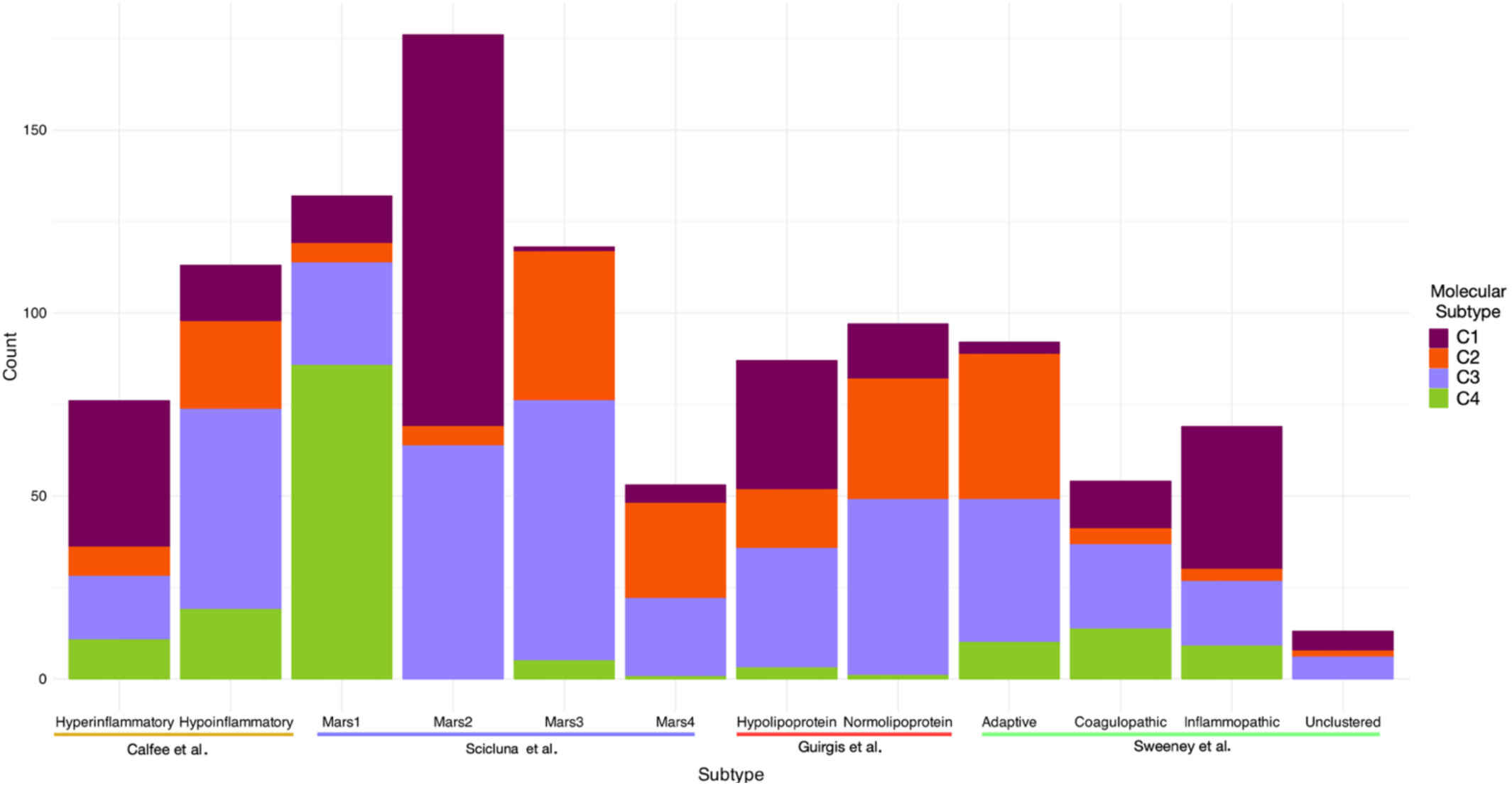
Molecular subtype enrichment compared to previous studies. Barplots showing the distribution of the molecular subtypes defined in this paper compared to subtypes from previous studies. Previous studies are listed on the x-axis and barplots are color-coded by our molecular subtypes.

**Supplemental Figure 3:**
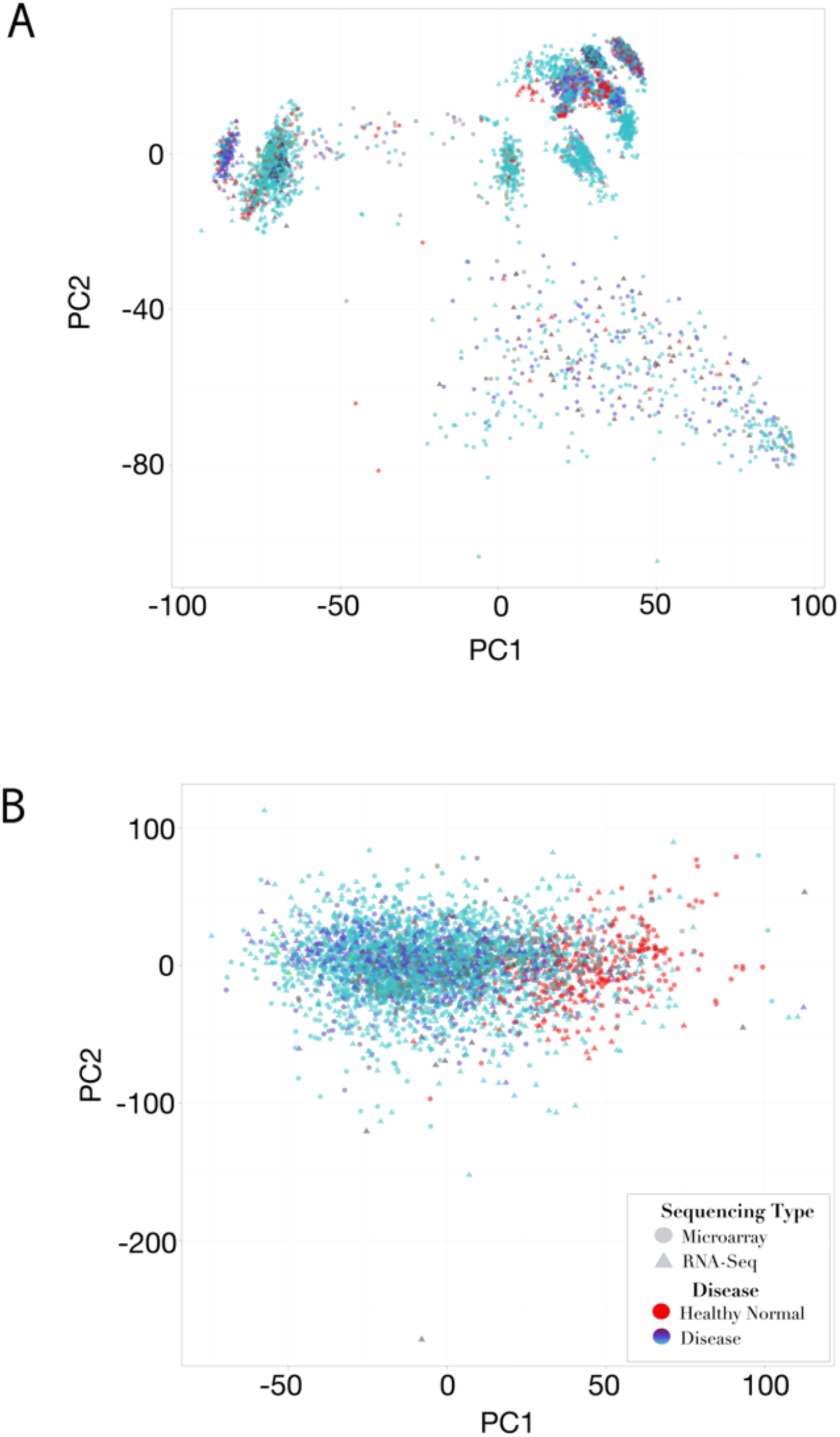
Samples before and after batch correction. (A) Samples from all datasets included in analysis in PCA space prior to performing batch correction. (B) Samples from all datasets included in analysis in PCA space after performing batch correction.

